## Supplementary for "Cognitive Reserve Against Alzheimer’s Pathology Is Linked to Brain Activity During Memory Formation"

### 5 Supplementary Material

#### 5.1 Supplementary Methods

##### 5.1.1 Quality control and sample cleaning

As another means to improve signal to noise ratio we chose strict criteria for outlier exclusion based on behavioral and task fMRI metrics. Individuals were excluded if either of the following was true: (1) They made more than 8 errors in their indoor/outdoor judgement. This corresponds to individuals with extreme outliers in the distribution of indoor/outdoor errors and could be related to lack of attention or confusion. (2) Based on response bias in their confidence rating during post-MRI retrieval, represented by the criterion location $c = -\frac{1}{2}(z(HR) + z(FAR))$; z = normal inverse cumulative distribution function, HR = hit rate, FAR = false alarm rate. Individuals with absolute response bias values above 1.5 were excluded, since strong bias could potentially render the parametric modulation invalid for two reasons. First, the response category would likely not correspond to the actual BOLD signal at the time of encoding. Second, a reliable estimation of the subsequent memory regressor does require some variability in the response categories. (3) Framewise displacement (FD) was above 0.5mm in a single EPI or above 0.2mm in more than 2% of the EPIs. This exclusion was supposed to limit motion effects on the data quality. (4) An individual had extreme outliers in the $\beta$ values of more than 10% of the voxels of their (GM-masked) regressor image. This was indicative of inaccurate estimations of the subsequent memory regressor in large parts of the brain and could have skewed the results of subsequent modeling steps. 68 individuals (11 CN, 6 ADR, 20 SCD, 19 MCI, 12 AD dementia) were excluded based on these criteria, leaving an fMRI sample with 490 individuals.

##### 5.1.2 One-dimensional pathological load score

Due to the nonlinearity of the disease progression trajectory along the AD continuum in 3D ATN space ($A\beta_{42:40}$ ratio, CSF p-tau, hippocampal volumes), a nonlinear dimensionality reduction method called t-SNE[46] was employed to reduce the dimension to one, yielding a single PL score per subject. Broadly speaking, t-SNE converts the Euclidian distances between datapoints in the high-dimensional space into conditional probabilities, which represent similarities. Likewise, conditional probabilities are defined for the low-dimensional counterparts. A perfect representation of the data in a lower-dimensional space would retain the conditional probabilities from the high-dimensional space between all pairs of datapoints. Hence, an optimal solution is sought by minimizing the mismatch between the conditional probabilities in both spaces, which is quantified via the Kullback-Leibler divergence. More details about the method can be found in Van der Maaten and Hinton[46]. Assuming that the biomarker progression profile across individuals is homogeneous, t-SNE would be able to extract this progression profile faithfully by retaining the similarities between datapoints from the ATN space as much as possible in the one-dimensional output space.

As the name suggests, t-SNE is a stochastic algorithm. Hence, the random seed was fixed to 617 to ensure reproducibility. The algorithm was applied to all 441 participants with complete ATN data available using a perplexity parameter of 50. One individual was considered an outlier and removed from all analyses, as visual inspection indicated that its assigned PL score was an erroneous representation of its AD biomarker status, as assessed from comparison to all other individuals' PL scores (see Fig. S3 for a plot including the declared outlier). Subsequently, the resulting score was scaled to fall into the range between 0 and 1 by subtracting the minimal value across participants and then dividing it by the maximal value. For reasons of interpretability, the scale was reversed such that increasing numbers of the PL score refer to increasing amounts of AD pathology.

Moreover, the algorithm was tested with five different choices of the perplexity parameter (10, 25, 30, 50, 100) and the results were correlated with each other to check stability. On average, the correlation was 0.950, indicating that the obtained score does not exhibit substantial dependence on the selected perplexity parameter. Robustness of the PL score was further checked by applying the t-SNE algorithm 1000 times, each time randomly holding out 10% of the data. The mean correlation over 1000 iterations was 0.945, suggesting a high robustness of the proposed PL score based on ATN.

###### 5.1.3 Cross-validation to determine optimal number of principal components

The optimal number of principal components $P$ was determined in a 10-fold cross-validation approach. Its results are shown in Fig. S5. Since not all participants with functional data also had CSF measures and thus a PL score, the data was stratified into two groups accordingly. This ensured similar proportions of both groups within each fold, as participants with missing values for any of the variables in the moderation model could still be used for principal component analysis (PCA). PCA was performed separately on the training data in each of the folds after mean-centering the masked functional (training) data. The coefficients of the moderation model from Eq. 2 were then determined for different numbers of principal components (1-25) using the least-squares method. With these coefficients the held-out (test) data was predicted and $R^2$ values between true and predicted PACC5 values (Box-Cox transformed) were calculated for each combination of fold and number of principal components. In order to ensure independence of a particular division into folds, this procedure was repeated 10 times with different partitioning of the data into folds. The optimal number of principal components was identified as the corresponding model with the highest mean $R^2$ value across the 10*10 predictions.

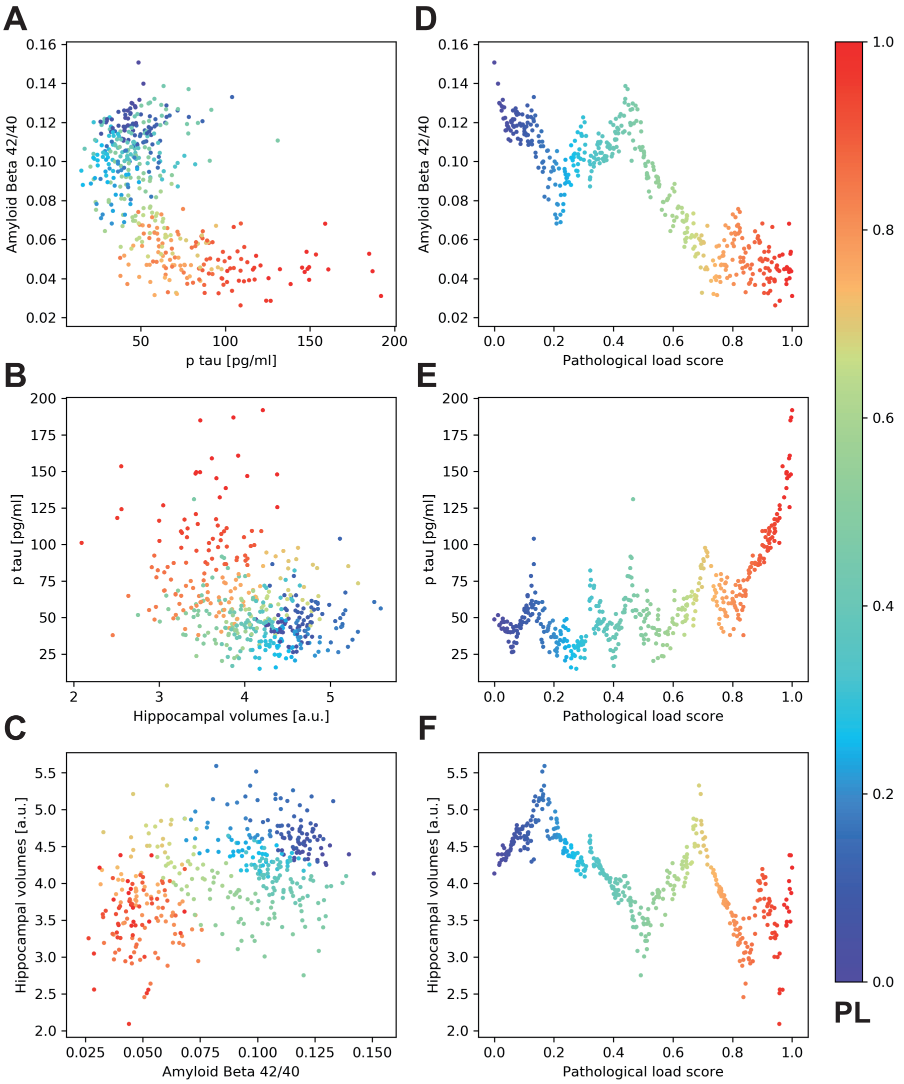


Figure S1: **Relationship between ATN biomarkers and data-driven PL score.** (A-C) All three possible combinations of pairs of biomarkers are shown. The points are color-coded based on the corresponding pathological load score. (D-F) Biomarkers plotted against the PL score. Hippocampal volumes refer to bilateral hippocampal volumes corrected for TIV. Please note that coloring of data points is redundant in panels D-F and was only done for illustrative purposes.

#### 5.2 Supplementary Results

##### 5.2.1 Construction of a continuous one-dimensional pathological load score

Based on the ATN framework that represents a biological characterization of AD,[45] a novel data-driven index for disease severity along the AD continuum was constructed combining CSF and MRI biomarkers of ATN in a one-dimensional score. The score ranges from 0 to 1, where 0 represents minimal AD pathology and 1 means maximal AD pathology in reference to the underlying sample. As demonstrated in Fig. S1, the underlying continuum follows a nonlinear pattern capturing dependencies across all three ATN biomarkers. More specifically, low $A\beta_{42:40}$ ratios (A), high p-tau measures (T), and small hippocampal volumes (N) result in high PL scores. Figs. S1A and C suggest that $A\beta_{42:40}$ is generally the strongest contributor to the PL score.

In the lowest range of PL scores from 0 to about 0.2, lower $A\beta_{42:40}$ ratios seem to be the main determinant of the PL score, which is thought to represent disease severity (Fig. S1). From roughly 0.2 until 0.5, higher values of the PL score are mainly characterized by lower hippocampal volumes (Fig. S1F). From there until PL scores of ca. 0.75 it is again primarily lower $A\beta_{42:40}$ ratios that contribute to a higher PL score. On the upper end of the PL score above 0.8, p-tau is the main dimension of variance (Fig. S1E).

Additionally, the correlation between the PL score and its single components was tested, to examine which dimension is (linearly) most strongly related to the PL score. The cross-correlations are -0.862 for $A\beta_{42:40}$, 0.670 for p-tau and $-0.684$ for the TIV-corrected hippocampal volumes. This confirms the impression that the A dimension had the biggest influence on the construction of the PL score. P-tau and the hippocampal volumes are reflected in the PL score to a similar extent, with different signs. This matches the observation that AD severity is associated with higher p-tau measures and lower hippocampal volumes.

In summary, the substantial correlations of the PL score with the ATN measures suggest shared variance and hence a meaningful reduction of the AD continuum to a single dimension.

##### 5.2.2 Pathological load is associated with cognitive performance

As a marker for disease severity on the AD continuum, the PL score was expected to show strong associations with the cognitive measures from neuropsychological testing. Indeed, we found a strong quadratic association between PL and PACC5 as measure of cognitive performance ($p = 8.78 \cdot 10^{-28}$, standardized regression coefficent $\beta$ = -0.501; Fig. 1) that was stronger than the linear association (see supplementary for a comparison). Indeed, using a linear model we found a strong association between PL and PACC5 as measure of cognitive performance ($p = 7.07 \cdot 10^{-22}$, standardized regression coefficent $\beta$ = -0.452). However, the empirical associations suggested rather a nonlinear relation, i.e. an increasing rate of cognitive decline with increasing pathology (Fig. 1). Hence, we further tested whether cognitive performance follows a quadratic function of the PL score. The association between (quadratic) PL and cognitive performance was significant ($p = 8.78 \cdot 10^{-28}$, $\beta$ = -0.501). In concordance with the differences in $\beta$, a comparison of the $R^2$ values reveals that the purely quadratic model fits the data better ($R^2 = 0.420$) than the linear model ($R^2 = 0.379$).

###### 5.2.3 CR score moderates effect of pathology on cognitive performance

Formal testing confirmed the interaction effect between the (quadratic) PL score and the CR score on PACC5 ($p = 4.64 \cdot 10^{-11}$, $\beta = 0.2713$) and the previously observed main effect of PL ($p = 1.45 \cdot 10^{-7}$, $\beta$ = -0.313).

##### 5.3 Supplementary Discussion

###### 5.3.1 PL score

A cautious note should be made about the PL score. One should be aware that it is a purely cross-sectional construct that is agnostic for the order of events along the disease progression towards Alzheimer's disease. It has to be stressed that biomarker levels at progressing PL scores hence cannot be interpreted as a sequence of events. In fact, according to the prevailing model of the Alzheimer's pathological cascade, abnormal levels of Aβ are the initiating event in AD,[49] followed by abnormal tau levels and neurodegeneration. However, we found that hippocampal volumes already showed a strong contribution to increasing PL scores at its lower levels, which would indicate atrophy as an early pathological change.

The challenges for a dimensionality reduction method in this context are many-fold. The t-SNE method used here tries to maintain neighborhood relationships between data points in lower-dimensional spaces. In a three-dimensional space, one could potentially find four data points that are mutually equidistant. There is no way to accurately retain these distances in a one-dimensional space. This is part of a challenge known as the "crowding problem".[46] Given the composition of the sample, which has an over-representation of individuals devoid of even early clinical symptoms, more severe AD biomarker profiles might not be represented faithfully in the PL score, as the algorithm tries to the preserve the distances beween the many less progressive AD biomarker profiles. On the other side, also age-related and other non-AD related pathological changes (e.g. hippocampal scleroris or vascular disease) might contribute to hippocampal atrophy, neurodegeneration and cognitive decline in our sample. This might be responsible for the stronger contribution of hippocampal volumes to lower PL scores. Furthermore, the influence of uncaptured covariates should not be underestimated. However, even if the sample uniformly represented individuals across different stages of the AD continuum, recent evidence for distinguishable AD subtypes (see e.g. Vogel et al.[50]) indicate the potential presence of multiple differential biomarker trajectories. In concert, these challenges might explain the unexpected pattern of trajectory reversals of the biomarkers with increasing PL scores that can be observed in Fig. S1.

Nevertheless, for the aim of examining cognitive reserve we make the simplified assumption that AD biomarkers can actually be represented by a single variable. The strong associations with the three biomarkers as well as with cognitive performance indeed suggest the PL score as a meaningful index of overall disease severity (rather than disease progression) for our purpose.

#### 5.4 Supplementary Figures

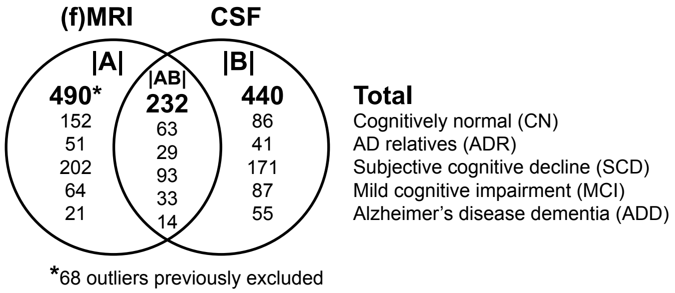


Figure S2: **Venn diagram of the sample**. Sample |A| refers to the participants with fMRI data and was used to derive eigen-images of the subsequent memory contrast images via PCA. Sample |B| refers to the participants with CSF data, which was used for creating the PL score. Sample |AB| is the union of both, i.e., the participants with both fMRI and CSF data. Sample |AB| was used to derive the CR-related activity patterns. The remaining participants of sample |B| that were not part of |AB| were used for further validation of the CR score by determining its ability to moderate the effect of neurodegeneration on individual measures of cognitive performance.

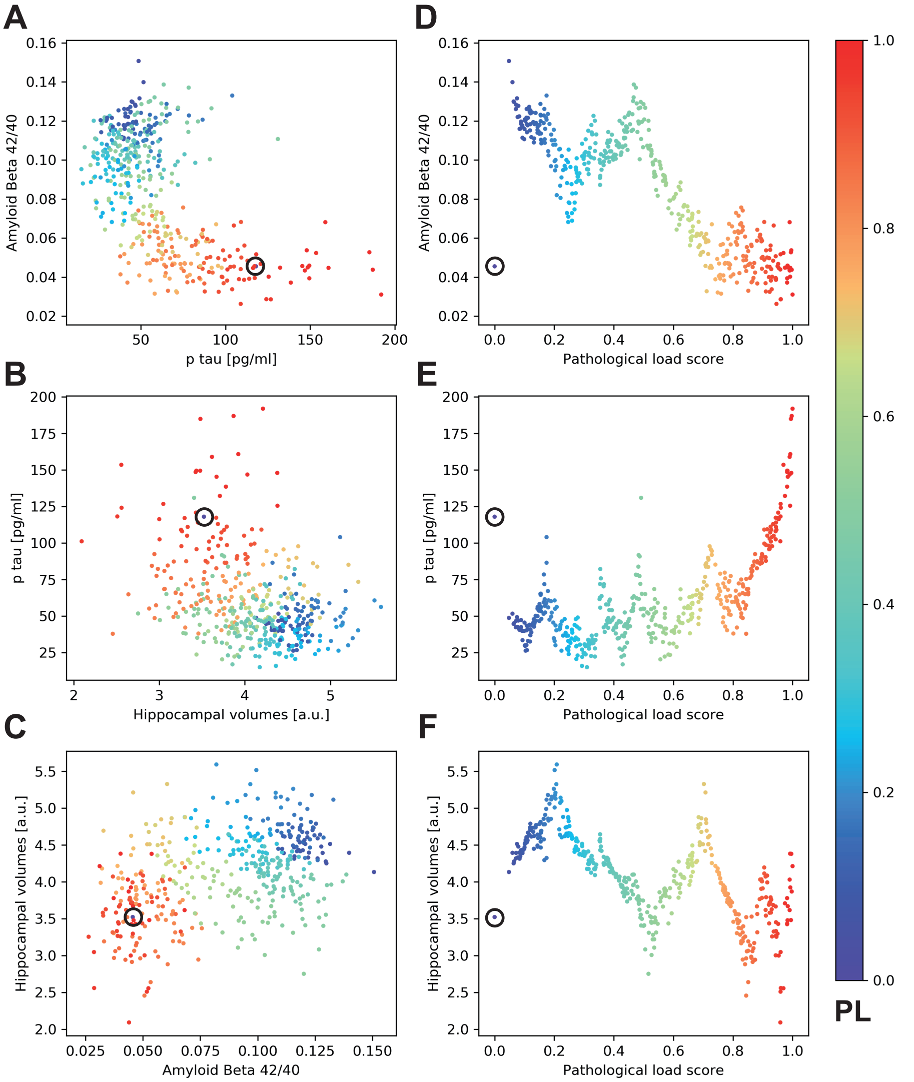


Figure S3: **Relationship between ATN biomarkers and data-driven PL score.** Same as Fig. S1, with the exception that one additional subject was included in the derivation of the PL score, which was deemed an outlier and excluded from all analyses. It is marked with a black circle and has a PL score of 0 despite its very low $A\beta_{42:40}$ ratio of 0.46, rather high p-tau levels of 118 pg/ml and TIV-corrected hippocampal volumes of 3.52.

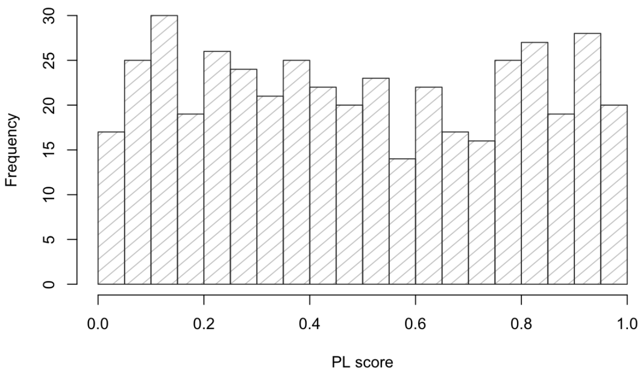


Figure S4: **Distribution of PL score**. The displayed sample is the full CSF sample (|B| in Fig. S2).

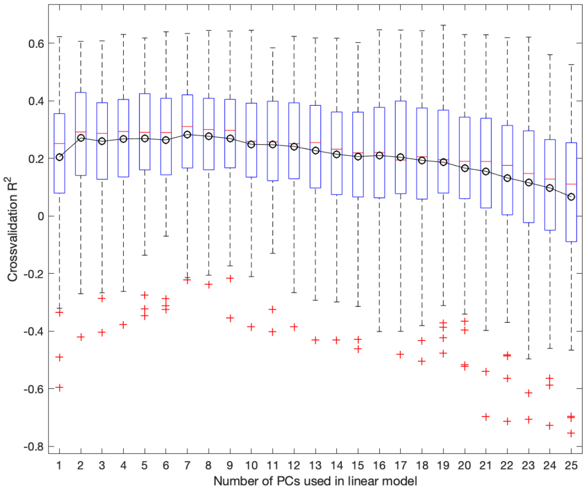


Figure S5: **Cross-validation results**. According to Eq. 2, PACC5 was predicted by varying numbers of principal components (eigen-images) in a 10-fold cross-validation procedure that was repeated 10 times with different partitioning of the data (see section 4.7 for details). The boxplots refer to the cross-validation $R^2$ in PACC5 scores (Box-Cox transformed) in the 10*10=100 independent test set predictions. The black line denotes the mean value across the 100 predictions.

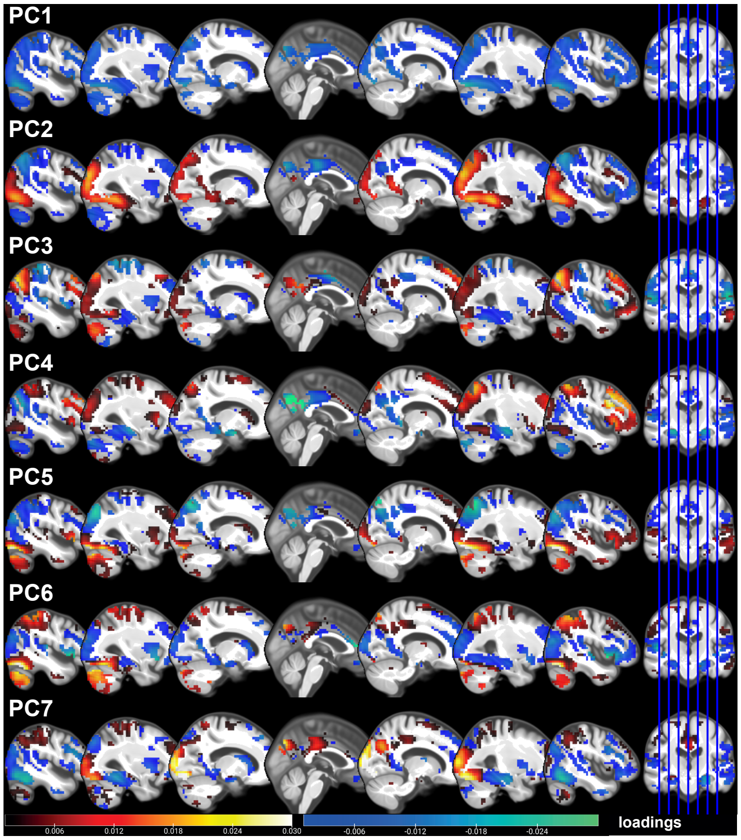


Figure S6: **Principal components of subsequent memory regressor images**. Shown are the loadings of each voxel onto each of the seven principal components (PC1-7) of the regressor images of subsequent memory. Please note that the PCA has been restricted to regions with a significant subsequent memory effect in the baseline sample (compare Fig. 2A) and hence does not include region outside of those 13695 voxels.

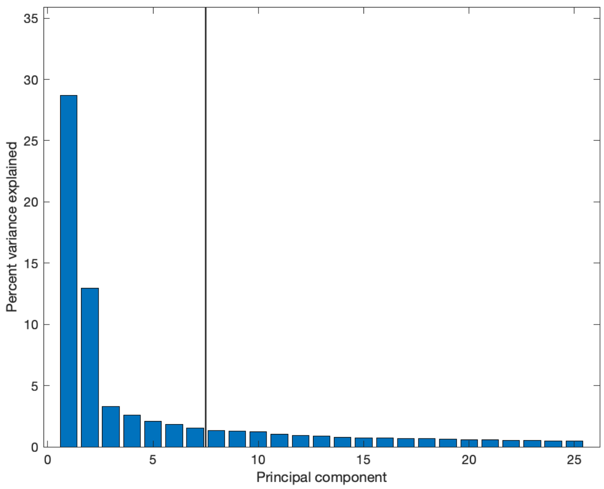


Figure S7: **Variance explained by PCs**. Shown is the amount of variance in the original subsequent memory regressor images that the individual principal components explain as assessed via their eigenvalues. The black line symbolises the optimal number of PCs determined via a 10-fold cross-validation procedure.
